# Temperature and sex shape reproductive barriers in a climate change hotspot

**DOI:** 10.1101/2023.02.12.528219

**Authors:** Cristóbal Gallegos, Kathryn A. Hodgins, Keyne Monro

**Affiliations:** School of Biological Sciences, Monash University, Melbourne, VIC, Australia

**Author notes:** Corresponding author: Cristóbal Gallegos.

**Keywords:** species barriers, hybridization experiment, hybrid fitness, extrinsic barriers, thermal performance, temperature-mediated barriers, maternal inheritance

## Abstract

Climate change is shifting species ranges and altering reproductive interactions within those ranges, offering closely-related species new scope to mate and potentially hybridize. Predicting hybridization and its outcomes requires assessing the interplay of biological and climatic factors that mediate reproductive barriers across life stages. However, few studies have done so across the range of environments that parents and offspring potentially encounter in nature, as is crucial to understand the environmental sensitivity of reproductive isolation and its fate under climate change. We set out to assess prezygotic and postzygotic reproductive barriers, and their dependence on temperature and sex, in sister species of a marine tubeworm (*Galeolaria*) from a sentinel region for climate change impacts in southern Australia. We performed reciprocal crosses within- and between-species using replicate populations, and assessed fertility of crosses, survival of embryos, and survival of larvae, at five temperatures spanning the thermal ranges of populations in nature. We found that barriers were weak and independent of temperature at fertilization, but stronger and more temperature-sensitive at larval development, as species diverged in thermal tolerance. Barriers were asymmetric between reciprocal hybrids, moreover, suggesting a complex interplay between thermal adaptation in parental lineages and maternal inheritance of factors (e.g., mitochondria, endosymbionts) that influence hybrid viability across temperatures. Together, our findings provide new insights into the roles of temperature and sex in reproductive barriers across early life stages, and point to shifting strengths of reproductive isolation in future climates.

## Introduction

Climate change is redistributing biodiversity and altering reproductive interactions in existing ranges, offering new scope for closely-related species to mate and hybridize (Chown et al., 2015; Chunco, 2014; Pecl et al., 2017). Whether they can do so, and the consequences for population dynamics, adaptation, and reproductive isolation, are often unknown (Todesco et al., 2016; Vallejo-Marín & Hiscock, 2016). Predicting hybridization and the fates of species barriers in future climates is therefore a key challenge at the interface of ecology, evolution, and conservation, and requires understanding the strengths of barriers as well as the biological and environmental factors that mediate them (Chunco, 2014; Vallejo-Marín & Hiscock, 2016). Doing so can provide new insights into the nature of reproductive isolation and its environmental sensitivity, with implications for proactive management and conservation of biodiversity under climate change (Genovart, 2008; Taylor et al., 2015).

Hybridization has diverse consequences that may be adverse or beneficial, and highly context-dependent (Todesco et al., 2016). On the one hand, hybridization can increase extinction risk if relatively fitter hybrids replace one or both parent species (genetic swamping), or if population growth rates decline due to wasteful production of maladapted hybrids (demographic swamping; Todesco et al. 2016). On the other hand, hybridization can enhance population growth and adaptive capacity by increasing heterozygosity, catalysing formation of novel phenotypes, or transferring adaptive alleles between species (Leroy et al., 2020; Rieseberg et al., 2003), though reproductive barriers may weaken as a result (Owens & Samuk, 2020). Such outcomes rest on the production and fitness of hybrid offspring, which can in turn depend on the divergence of parental lineages (Edmands, 2002; MacPherson et al., 2022) and the selection regimes that parents and offspring face (Arnold & Martin, 2010). Since climate change will likely alter selection (Hoffmann & Sgrò, 2011; Siepielski et al., 2017), a better understanding of climatic impacts on reproductive barriers, and insights from regions where reproductive interactions are exposed to rapid change, are urgently needed.

Reproductive barriers are classed as prezygotic or postzygotic, based on when they manifest (Coughlan & Matute, 2020; Coyne & Orr, 2004). Prezygotic barriers prevent hybrids from forming by deterring either heterospecific matings (e.g., through reproductive asynchrony, or mismatched genitalia; Mangubhai & Harrison, 2008; Sánchez-Guillén et al., 2014) or heterospecific fertilizations once gametes interact (e.g., through gamete recognition systems; Kosman & Levitan, 2014). Postzygotic barriers, in contrast, prevent hybrid survival and reproduction (Fierst & Hansen, 2010), and frequently result from genetic incompatibilities between parental lineages (Coyne & Orr, 1989; Miller & Matute, 2017). They can emerge at different points during development, especially in complex life cycles spanning distinct stages that often differ in form, function, or duration (Albecker et al., 2021; Marshall et al., 2016). Nevertheless, the relative impacts of prezygotic and postzygotic barriers on reproductive isolation are unclear. Although postzygotic barriers dominated early work (Dobzhansky, 1937; Haldane, 1922), insights since then — for example, that prezygotic barriers are stronger, and evolve faster, in sympatric *versus* allopatric species to limit gamete wastage on hybrids (Coyne & Orr 1989, 1997) — have shaped the modern view that prezygotic barriers are more prevalent and influential (Coughlan & Matute, 2020; Lowry et al., 2008). Ultimately, understanding when barriers arise, and their strengths at different life stages, is a key step in inferring isolation and its underpinnings (Bundus et al., 2015).

Reproductive barriers are further classed as intrinsic or extrinsic, based on their environmental sensitivity (Coughlan & Matute, 2020). Either type can emerge before or after zygote formation, but intrinsic barriers are considered robust to environmental context (per the examples above), whereas extrinsic barriers rely on environmental factors for their maintenance. Temperature, for example, maintains prezygotic barriers between fruit fly species by shaping differences in thermal niche (Matute et al., 2009) or morphological plasticity in genitalia (Peluffo et al., 2021). The distinction between intrinsic and extrinsic barriers can be blurred, however, if strengths of intrinsic barriers vary environmentally. For instance, hybrid incompatibilities (encompassing inviability and sterility) can strengthen or weaken depending on the temperatures experienced as offspring develop (Miller & Matute, 2017), and those to which parental lineages are adapted (Keller & Seehausen, 2012). Hybrids can also tolerate stressful environments better than parental lineages, which risk being displaced in such cases (Hwang et al., 2016; Martins et al., 2019). Assessments of reproductive barriers across the range of environments that parents and offspring may face are therefore vital for understanding the environmental sensitivity of reproductive isolation, and predicting its response to future environments, but are currently lacking (Coughlan & Matute, 2020; Martins et al., 2019).

Last, reproductive barriers may depend on sex in ways that interface with the barriers above. Haldane’s rule poses that hybrids of the heterogametic sex tend to have lower fitness than those of the homogametic sex (Haldane, 1922) due to differences in chromosomal dynamics or sexual selection (Panhuis et al., 2001; Turelli & Orr, 1995; Wu & Davis, 1993). A related phenomenon, dubbed Darwin’s corollary to Haldane’s rule (Turelli & Moyle, 2007), is asymmetry in the fitness of reciprocal hybrids (those derived from sperm *versus* eggs of parental lineages) due to incompatibilities involving uniparentally inherited factors, such as organelles or endosymbionts (Bruzzese et al., 2022; Gebiola et al., 2016). Despite the potential for environmental conditions to mediate Haldane’s rule (Bundus et al., 2015; Wade et al., 1999), evidence of how the environment could mediate its corollary seems scarce (Falk et al., 2012). Consequently, we have limited understanding of the interplay between parent-of-origin effects and external environments, calling for new insights into hybrid production and fitness under diverse environmental conditions (Coughlan & Matute, 2020), with those relating to climate change especially.

Here, we assess the effects of temperature and sex on barriers to reproduction in sister foundation species from southern Australia (Figure 1), a global marine hotspot harbouring exceptional biodiversity and warming faster than the planetary average (Costello et al., 2022; Hobday & Pecl, 2014). The tubeworm *Galeolaria* has a complex life cycle with early life stages (gametes, embryos, and larvae) that fertilize and develop in the external environment (Chirgwin et al., 2021; Rebolledo et al., 2020), offering rare power to isolate the role of temperature in reproductive isolation at different stages. Sister species are sympatric and breed synchronously in the hotspot, but share little gene flow (Gallegos et al., 2023). That they also show genomic signals of thermal adaptation (Gallegos et al., 2023), and *G. caespitosa* is restricted to higher latitudes whereas *G. gemineoa* extends almost to the tropics (Halt et al. 2009), suggests that thermal niche differentiation could be involved in their reproductive isolation. We therefore set out to test whether barriers between species differ in strength depending on life stage and parent of origin, and how they are affected by potential differences in thermal niche. To do so, we crossed six replicate populations per species in a split-cohort design with genetic and environmental backgrounds standardized across life stages, and assessed the fertility of crosses, survival of embryos, and survival of larvae, across five temperatures spanning species’ thermal ranges. We performed crosses reciprocally to assess parent-of-origin asymmetry in barriers, and within species to assess thermal niches at each stage. Our findings shed new light on the roles of temperature and sex in reproductive barriers across early life stages, and point to shifting strengths of isolation in future climates.

**Figure 1.**
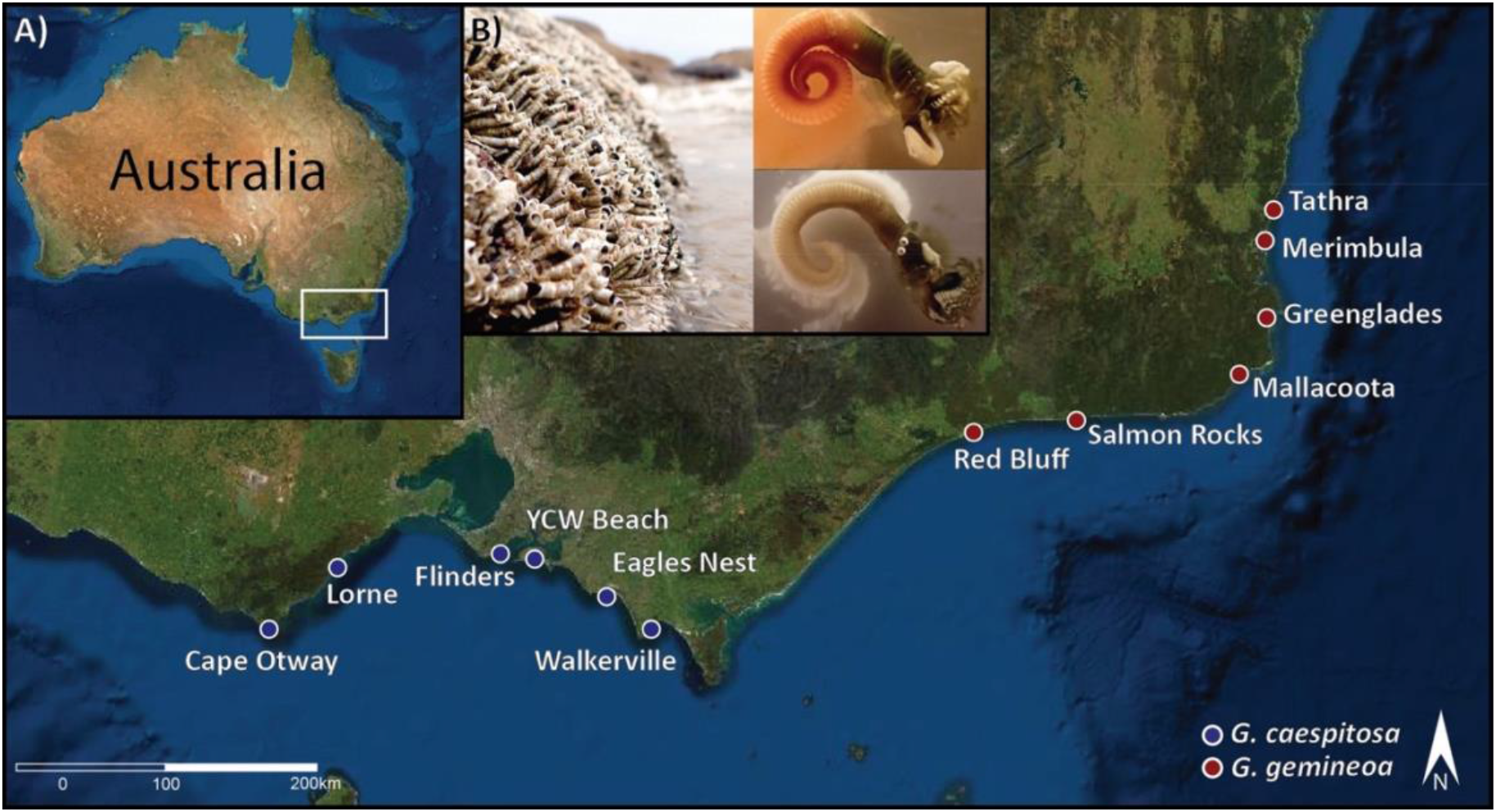
Sampling of the tubeworms *Galeolaria caespitosa* (blue points) and *G. gemineoa* (red points) in the southern Australian hotspot (inset A), a region of exceptional marine biodiversity and accelerated warming. Inset B shows a typical colony with adults retracted into tubes at low tide (left), and different sexes releasing eggs (top right) or sperm (bottom right) after extraction from tubes.

## Methods

### Study system and sampling

*Galeolaria* tubeworms (Serpulidae) are cryptic foundation species on rocky shores of southern Australia, where their dense colonies of tubes enhance biodiversity by forming habitat for species that cannot otherwise persist there (Figure 1B; Wright & Gribben, 2017). The nature of sex determination is unknown, but mature adults are differentiated by the presence of sperm or eggs (Figure 1B) that they shed year-round into the sea to fertilize externally (Chirgwin et al., 2020; 2021). Embryos develop into functionally independent larvae ∼24-72 hours later, and larvae spend another ∼2–3 weeks feeding and growing while dispersed by currents, before transitioning to sessile life stages (juveniles and adults) onshore (Marsden & Anderson, 1981; Rebolledo et al., 2020). As for other aquatic ectotherms, these early pelagic stages are thermal bottlenecks in the lifecycle, defining population abundances and genetic structures across species’ ranges (Dahlke et al., 2020; Lotterhos et al., 2021; Rebolledo et al., 2020). These stages also differ markedly in form, function, and duration (Byrne, 2011; Rebolledo et al., 2020), offering different scope for barriers arising during hybrid production and development to interact with environmental temperature.

We sampled six populations per species across the hotspot (Figure 1) between November 2020 and June 2021, sampling each block of populations in our experimental design (detailed below) in the same week (Table S1). Sympatric populations were avoided, given the challenge of differentiating sister species in the field (Gallegos et al., 2023; Halt et al., 2009). Adults were kept isolated by population and transported in aerated coolers to Monash University within days of collection. There, they were housed by population in aerated aquaria at 17 °C with water refreshed as needed, and fed a mix of live microalgae twice weekly. Since gametogenesis is seemingly continuous, and gametes can ripen in under two weeks (Chirgwin et al., 2017), adults were acclimated to these conditions for three weeks to reduce environmental variation before collecting gametes for crosses. Remaining population-level variation among crosses is therefore assumed to be mostly genetic in origin.

### Experimental design for pure and hybrid crosses across temperatures and life stages

Crosses were performed in blocks given we could not perform them all at once. Each block comprised two populations per species, crossed as in Figure 2 to give two pure (within-population) crosses of *G. caespitosa*, two pure (within-population) crosses of *G. gemineoa*, and eight hybrid (between-species) crosses. Hybrid crosses were four unique interspecific combinations that were each crossed reciprocally (i.e., *G. caespitosa × G. gemineoa* and *G. gemineoa × G. caespitosa*, where species listed first contributed eggs and species listed second contributed sperm; Figure 2). Between-population crosses were less relevant to our aims and were therefore omitted for feasibility. Each cross was initiated using sperm from 10 males and eggs from 10 females, and was replicated twice at each of five temperatures (12, 15, 17, 22, and 25 °C) at each of three life stages (fertilization, embryogenesis, and larval development), as detailed below. Temperatures spanned the ranges experienced by early life stages in nature, averaging ∼12–17 °C in winter and ∼19–25 °C in summer from southwest to northeast (www.ghrsst.org; Frusher et al., 2014; Stobart et al., 2015), and maintained within 0.1 °C using mini dry baths (Benchmark Scientific) for fertilization and embryogenesis or immersion heaters (Grant Optima TX150) for larval development.

**Figure 2.**
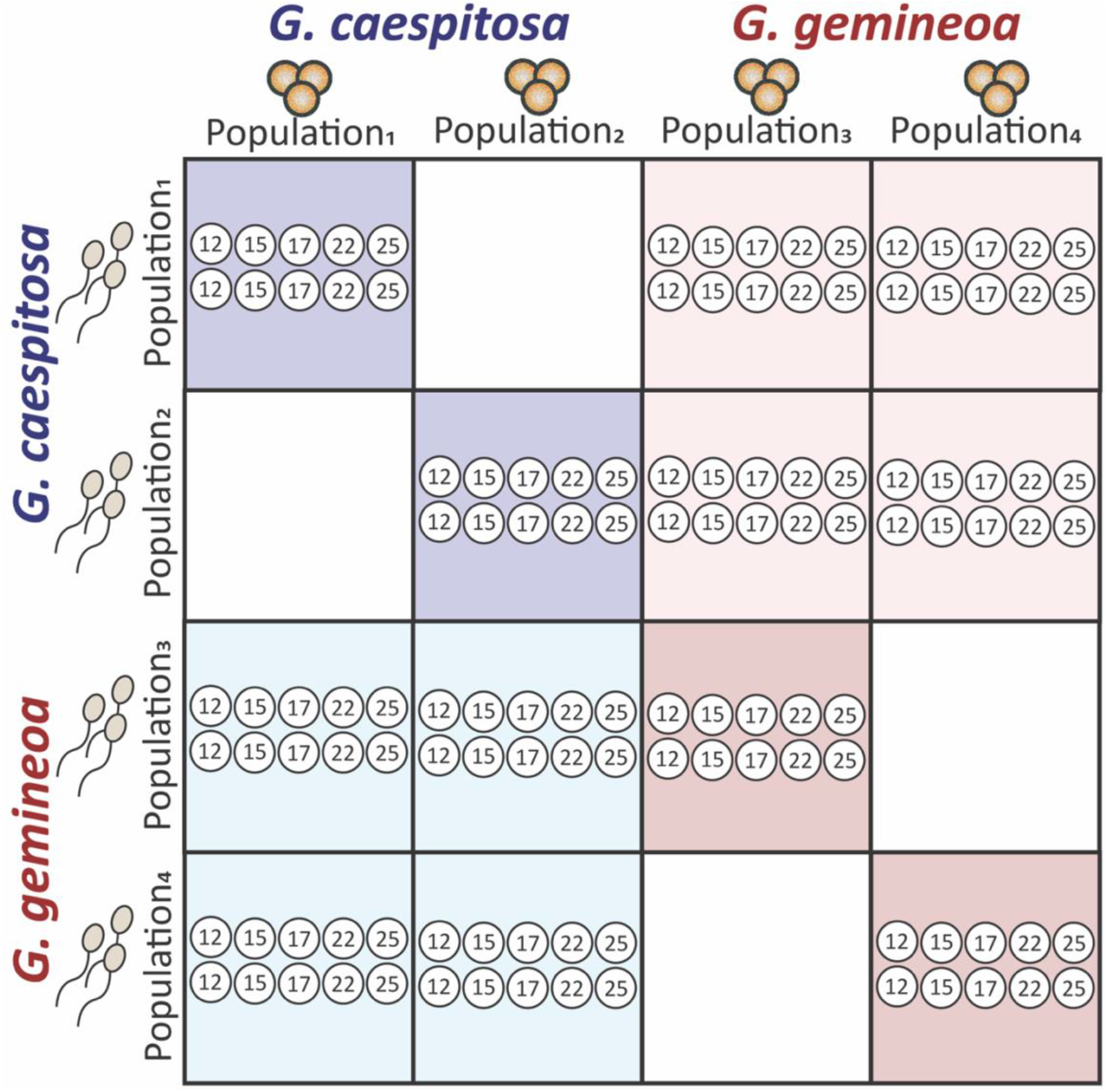
Experimental design for pure (within-population) and hybrid (between-species) crosses across temperatures (12, 15, 17, 22, and 25 °C) and early life stages (fertilisation, embryogenesis, and larval development, omitted here for simplicity). Crosses were performed in blocks of two populations per species (with *G. caespitosa* in blue and *G. gemineoa* in red), sampled from locations in Figure 1. Each cross was initiated using sperm pooled from 10 males (rows) and eggs pooled from 10 females (columns), and was replicated twice per temperature, parent of origin, and stage.

The design had three such blocks in total, with the first block repeated for all life stages, and the third block repeated for larval development, to compensate for mortality. By the end of the experiment, this produced 12 pure crosses and 24 hybrid crosses, with ∼60,000 fertilizations, embryos, or larvae scored across nearly 1,500 replicates overall (36 crosses × 5 temperatures × 3 life stages × 2 replicate vials per combination, plus repeated blocks).

### Gamete collection and fertilization protocol

For each block of crosses in Figure 2, gametes were collected by extracting each adult from its tube and placing it in minimal (200 μL) filtered sea water at 17 °C to spawn. After spawning, 100 μL of sperm or eggs per adult were pooled by sex in two separate vials, with concentrations in vials estimated from replicate cell counts in a hemocytometer. Since sperm are activated by dilution (Kupriyanova & Havenhand, 2002), gametes were kept undiluted at ∼4 °C to minimise aging until all adults had been spawned (∼2-5 hours). Pilot work showed that this protocol produced similar fertilization rates as when gametes were crossed sooner after spawning.

In preparation for crosses, eggs from each population were diluted with filtered, pasteurized seawater to ∼2,000–2,500 cells/mL, then 900 μL of eggs were pipetted into each of 30 replicate vials and adjusted to their test temperatures for ∼15 minutes. Sperm from each population were likewise diluted to ∼10^8^ cells/mL and adjusted to their test temperatures for ∼5 minutes. Each cross was then initiated by pipetting 100 μL of sperm per population into a corresponding vial of eggs, and mixing the solution by gentle inversion. The final sperm concentration (10^7^ cells/mL in a total volume of 1 mL) maximized fertilization rates in pilot work, and oxygen limitation is unlikely at this volume given the small sizes of gametes (Chirgwin et al., 2018).

After one hour of contact (sufficient for fertilisation rates to plateau at each temperature; Rebolledo et al., 2020), sperm were removed to further minimize the risks of oxygen limitation and polyspermy. This was done by carefully pipetting out most of the solution above eggs (which sink to the bottoms of vials), replacing it with 1 mL of fresh filtered seawater at the same temperature, then repeating the procedure 10 minutes later once eggs had re-sunk.

### Scoring fertilization success and offspring survival in crosses

#### Fertilization success

Vials of crosses used to score fertilization success were preserved by adding N0.1 mL of Lugol’s solution 1.5–4.5 hours after sperm removal, depending on temperature. Fifty eggs were then sampled randomly from each vial, examined under a microscope, and scored as fertilized if they had a fertilization envelope (including a raised cone at the site of sperm fusion) or begun to cleave, or as unfertilized if they had not. Since the envelope and cone form minutes after fertilization in *Galeolaria* (Marshall & Bolton, 2007), eggs were preserved and examined after sufficient time to reliably detect fertilization (if present) at even the coldest test temperature.

#### Survival of embryogenesis

For vials of crosses used to score survival of embryogenesis, effects of temperature were isolated by performing crosses at 17 °C and maintaining them at 17 °C for another 2.5 hours to complete initial cleavages. Then, 50 embryos with 2–4 cells each (ensuring a similar point in development) were transferred from each vial to a new vial with 1 mL of filtered, pasteurized seawater and adjusted to their test temperature. Embryos were scored for successful completion of embryogenesis (development into actively swimming larvae) after 24 hours at 22– 25 °C, 48 hours at 17 °C, and 72 hours at 12–15 °C. Embryos that had died, or not yet completed development, were scored as unsuccessful. Based on previous work (Rebolledo et al., 2020; Belcher et al., in prep.), embryos that do not complete development by these times at these temperatures never do so.

#### Survival of larval development

For vials of crosses used to score survival of larval development, effects of temperature were isolated by performing all crosses at 17 °C and maintaining them at 17 °C for another 48 hours to complete embryogenesis. Then, 30 actively swimming larvae were transferred from each vial to a new vial with 5 mL of filtered, pasteurized seawater, adjusted to their test temperature, and fed a mix of live microalgae *ad libitum* (∼10^8^ cells per vial thrice weekly). Larvae were scored for successful completion of development (changes in size, morphology, and behaviour signalling onset of metamorphosis; Marsden & Anderson 1981; Nelson et al., 2017) after 13 days at 25 °C, 15 days at 22 °C, 17 days at 17 °C, 20 days at 15 °C, and 24 days at 12 °C. Development times were based on larvae sampled from vials reserved for monitoring, without disturbing larvae in focal vials (focal larvae were scored only when monitored larvae had completed development or died), and matched previous work (Rebolledo et al., 2020).

### Data analyses

#### Estimating thermal tolerance curves

We analysed binary data (with 1 denoting successful fertilization or survival, and 0 denoting lack of success) using binomial mixed-effects regression models fitted with Laplace approximation in the *lme4* package (v1.1-27.1; Bates et al., 2015) for R (v4.0.5. R Core Team, 2021). Specific models are described below. Success was modelled as a quadratic function of temperature to estimate thermal tolerance curves for groups. Although classic descriptors, such as maximum, optimum (temperature at maximum), and breadth (temperature range at half maximum; Sinclair et al., 2016), were not formally estimated, temperature was centred and standardised to make model coefficients similarly interpretable (Schielzeth, 2010).

Each curve had an intercept estimating mean success, a linear trend estimating mean slope, and a quadratic trend estimating curvature. Curves were always concave down if quadratic trends differed from zero, and were broader (or narrower) if those trends were weaker (or stronger).

Concave curves had optima at the mean temperature if linear trends did not differ from zero, and had relatively higher (or lower) optima if those trends were positive (or negative).

#### Comparing prezygotic and postzygotic thermal tolerance among crosses

To compare thermal tolerance at different life stages among crosses, we fitted a model with cross, stage, and all possible interactions with linear and quadratic trends as fixed effects. Block (five groups), vial (1395 groups), and cross identity (36 groups based on unique combinations of population and parent of origin) were modelled as random effects. Random effects were usually intercepts, but linear trends also varied among cross identities to improve model fit (*χ*^*2*^ = 10.28, *d*.*f*. = 2, *P* < 0.01). Corresponding quadratic trends failed to improve fit (*χ*^*2*^ = 1.83, *d*.*f*. = 3, *P* = 0.61) and were omitted. More complex random effects (trends grouped by pure *versus* hybrid crosses) also failed to improve fit (*χ*^*2*^ = 6.33, *d*.*f*. = 3, *P* = 0.09) and were omitted.

Checks of model assumptions using the *DHARMa* package (version 0.4.4; Hartig, 2021) detected no violations. Significance of fixed effects was tested using Wald *χ* ^*2*^ tests (Bolker et al., 2009) in the *car* package (version 3.0-11; Fox & Weisberg, 2019). Groups in significant fixed effects were compared using pairwise contrasts (adjusted for multiple testing by controlling the false discovery rate) in the *emmeans* package (version 1.7.0; Lenth et al., 2021), and contrasts were visualised using its *emmip* function. Significance of random effects was tested using *χ* ^*2*^ likelihood ratio tests of nested models with *versus* without the effects of interest (Bolker et al., 2009).

#### Population-level variation in thermal tolerance

Since the model above suggested that thermal tolerance curves vary among populations in ways that could support local adaptation, we also explored population-level variation in tolerance. To do so, we re-fitted the model to success at each life stage for pure (within-species) crosses only. In this case, we modelled random intercepts and trends (as uncorrelated given the modest numbers of populations), and explored nested models with *versus* without random effects of interest grouped by species to compare variation between species and characterise it within species. To visualise variation, we predicted population-level tolerance curves from the model, together with their confidence intervals, using the *merTools* package (version 0.5.2; Knowles & Frederick, 2020).

## Results

### Differences in prezygotic and postzygotic thermal tolerance among crosses

#### Overview

We detected significant, interactive effects of cross and life stage on tolerance curves relating success (in terms of fertilization or postzygotic survival) to temperature (Table 1). The interaction between cross and life stage indicated that differences in mean success among crosses depended on stage (Figure 3). Likewise, the interaction between cross, life stage, and linear trend for temperature indicated that differences in thermal optima among crosses also depended on stage (Figure 3). Although there were significant interactions involving the quadratic trend for temperature (Table 1), pairwise contrasts (Figure 4G-I) indicate these were due to differences in thermal breadth among stages but not among crosses within stages, with curves effectively linear for fertilisation success (Figure 3A), but significantly concave for postzygotic survival (Figure 3B-C).

**Table 1.** Effects of cross (pure species or hybrids) and life stage (fertilization, embryogenesis, or larval development) on thermal tolerance, measured by success in terms of fertilization or postzygotic survival at different test temperatures. Success was modelled as a quadratic function of temperature in a binomial mixed-effects regression. Tolerance curves predicted by the model are shown in Figure 3, and contrasts of curve elements (means, linear trends, and quadratic trends) among groups are shown in Figure 4.

| Fixed effects | $\chi^2$ | d.f. | *P*-value |
| --- | --- | --- | --- |
| Cross | 11.79 | 3 | < 0.01 |
| Life stage | 73.60 | 2 | < 0.001 |
| Temperature (linear trend) | 44.63 | 1 | < 0.001 |
| Temperature (quadratic trend) | 107.56 | 1 | < 0.001 |
| Cross × life stage | 52.64 | 6 | < 0.001 |
| Cross × temperature (linear trend) | 37.70 | 3 | < 0.001 |
| Cross × temperature (quadratic trend) | 7.61 | 3 | 0.05 |
| Life stage × temperature (linear trend) | 2.98 | 2 | 0.23 |
| Life stage × temperature (quadratic trend) | 37.82 | 2 | < 0.001 |
| Cross × life stage × temperature (linear trend) | 32.89 | 6 | < 0.001 |
| Cross × life stage × temperature (quadratic trend) | 4.00 | 6 | 0.68 |

**Figure 3.**
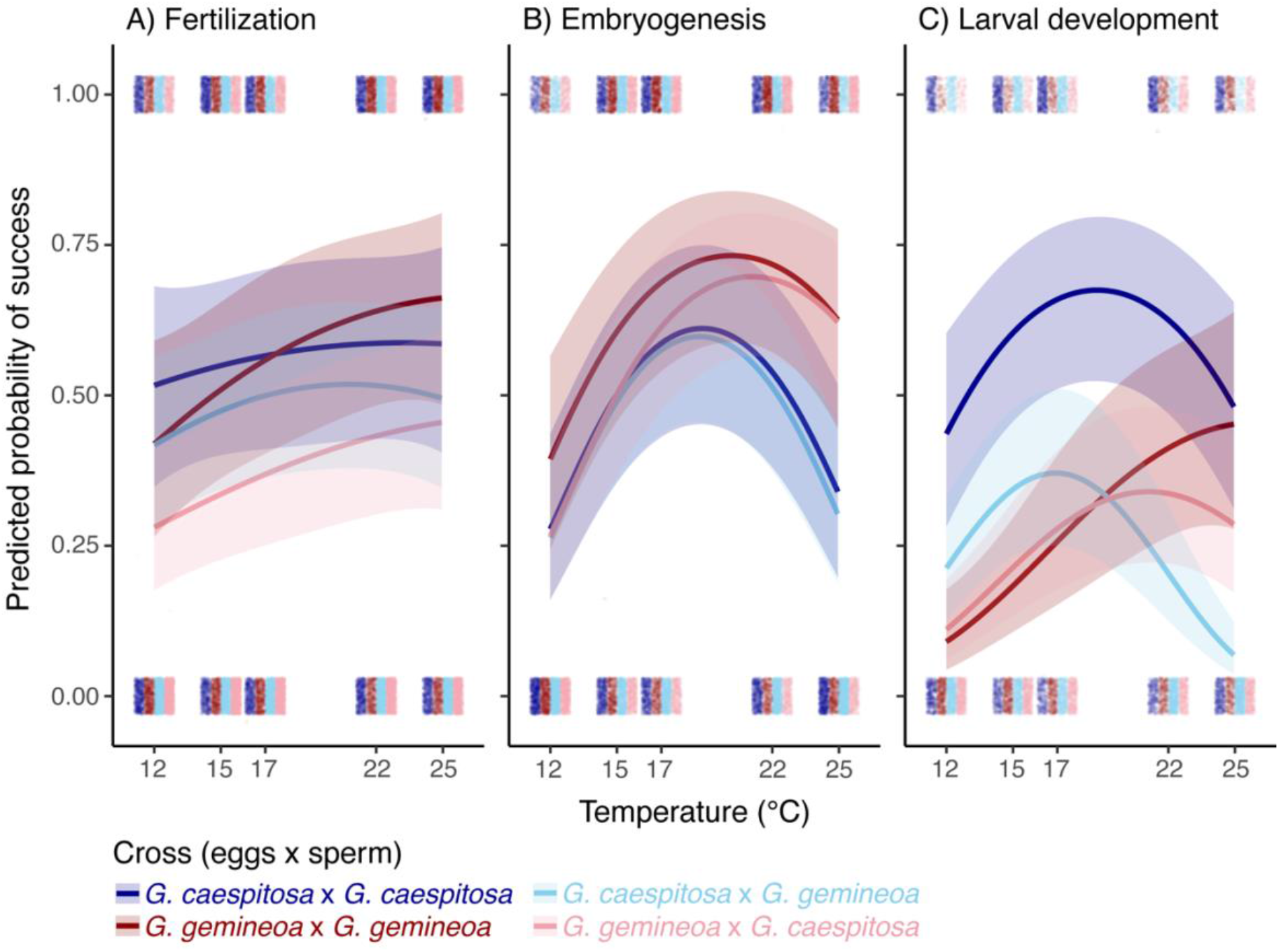
Thermal tolerance curves showing the predicted probabilities of A) fertilization success, B) survival of embryogenesis, and C) survival of larval development for pure and hybrid crosses. In hybrid crosses, species listed first contributed eggs and species listed second contributed sperm. Curves are predicted from a binomial mixed-effects regression of success (in terms of fertilisation or postzygotic survival) on temperature interacting with cross and life stage. Shaded areas indicate 95% confidence intervals of curve predictions. Points are raw data scored for each temperature, cross, and stage, with 1 denoting success and 0 denoting lack thereof. To aid visualization, points are transparent and spread both vertically and horizontally, so that darker colours indicate higher densities.

**Figure 4.**
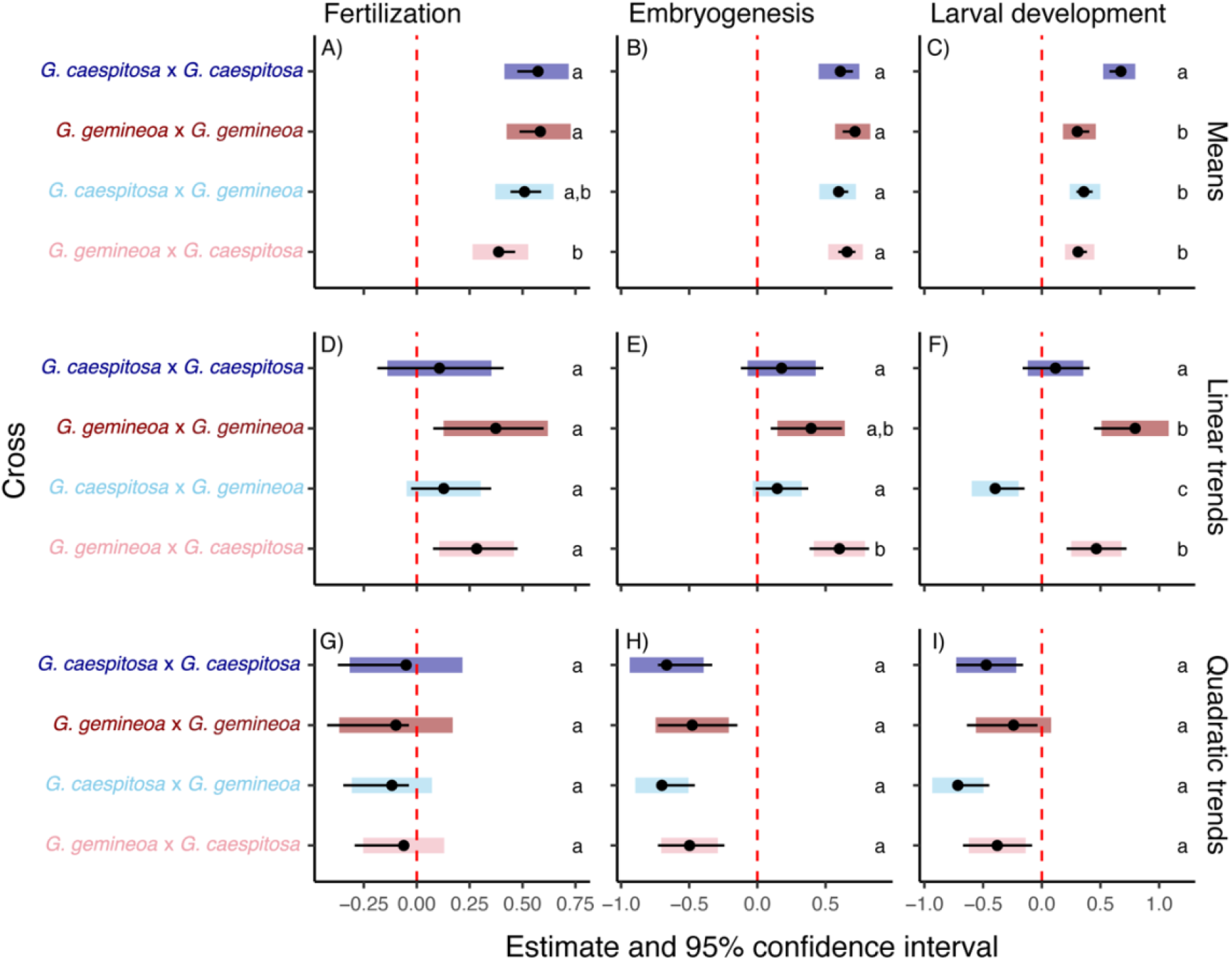
Means (A-C), linear trends (D-F), and quadratic trends (G-I) for thermal tolerance curves in Figure 3, contrasted between pure and hybrid crosses. Means and trends differ significantly from 0 if their estimates (points) have 95% confidence intervals (coloured bars) excluding zero, marked by red dashed lines. Means and trends differ significantly between crosses if 95% confidence intervals of their pairwise contrasts (black lines) do not overlap. Differences between crosses are also summarised by letters at the right of each panel (crosses sharing a letter do not differ from each other at *α* = 0.05).

#### Pairwise contrasts of crosses at different life stages

Species barriers at fertilization differed between parents of origin, with mean fertility being similar for pure crosses and hybrid crosses with *G. gemineoa* sperm, but significantly lower for hybrid crosses with *G. gemineoa* eggs (Figure 3A and 4A). Barriers at this stage depended little on temperature, since linear trends relating fertility to temperature did not differ among crosses, despite being positive for crosses with *G. gemineoa* eggs and nonsignificant otherwise (Figure 3A and 4D). Quadratic trends relating fertility to temperature were nonsignificant (Figure 3A and 4G).

Barriers at embryogenesis were generally weak, with all crosses having similar mean survival at this stage (Figure 3B and 4B). However, linear trends relating survival to temperature now differed among crosses, being significantly steeper (indicating a higher thermal optimum) for maternal crosses of *G. gemineoa* compared to other crosses (Figure 3B and 4E). Quadratic trends were negative and similar in magnitude (indicating similar thermal breadth) for all crosses (Figure 3B and 4H).

Last, species barriers grew stronger and more temperature-dependent at larval development.

Mean survival at this stage was higher for pure crosses of *G. caespitosa* compared to other crosses (Figure 3C and 4C), and linear trends relating survival to temperature differed between pure crosses, pure and hybrid crosses with *G. caespitosa* mothers, and reciprocal hybrids (Figure 3C and 4F). Specifically, linear trends were positive for all crosses with *G. gemineoa* mothers, negative for hybrid crosses with *G. caespitosa* mothers, and nonsignificant for pure crosses of *G. caespitosa*. Quadratic trends were negative except for pure crosses of *G. gemineoa*, whose nonsignificant trend suggested a thermal optimum above 25 °C, and were similar in magnitude among crosses (Figure 3C and 4I). Together, results for this stage supported three conclusions.

First, *G. gemineoa* (the northern species) had a higher thermal optimum than *G. caespitosa* (the southern one). Second, thermal optima differed among crosses in ways that weakened barriers at some temperatures (i.e., ∼19 °C, where hybrids and pure crosses of *G. gemineoa* survived similarly well) but strengthened them at others (i.e., above ∼19 °C, where pure crosses increasingly converged in survival and hybrids increasingly died; Figure 3C). Third, thermal optima differed between reciprocal hybrids in ways that suggest control of thermal niche by mothers (i.e., hybrid optima approximated those of maternal lineages; Figure 3).

#### Population-level variation in thermal tolerance

Although random effects in our main model supported variation in thermal optima among cross identities, this was unrelated to variation among populations (pure crosses; Figure S1). Population-level variation did not differ between species (comparison of models with *versus* without random effects grouped by species: *χ*^*2*^ = 0.01, *d*.*f*. = 3, *P* = 0.99), nor in relation to temperature (comparison of models with *versus* without random trends: *χ*^*2*^ = 0.01, *d*.*f*. = 2, *P* = 0.99). Rather, populations varied only in mean success (comparison of models with *versus* without random intercepts: *χ*^*2*^ = 7.08, *d*.*f*. = 1, *P* < 0.01), thus suggesting no differences in thermal optima and offering little support for local thermal adaptation in the phenotypes or locations sampled.

## Discussion

As climate change redistributes Earth’s biodiversity, predicting the outcomes for reproductive interactions between species requires understanding species barriers and their dependence on climate (Chunco, 2014; Vallejo-Marín & Hiscock, 2016). Here, we assessed prezygotic and postzygotic barriers across early life stages, their sensitivity to temperature, and their sexual asymmetry, in sister *Galeolaria* species whose ranges overlap in a sentinel region for climate change impacts (Gallegos et al., 2023). Barriers were strongest and most temperature-sensitive during larval development, and suggest a complex interplay between adaptation to different thermal niches and maternal inheritance. Specifically, larvae of the northern species (*G. gemineoa*) had a higher thermal optimum for survival than larvae of the southern species (*G. caespitosa*), and maternal hybrids of *G. gemineoa* survived better at higher temperatures than maternal hybrids of *G. caespitosa*. Maternal hybrids of *G. gemineoa* may be rarer, however, given that hybrid crosses initiated with *G. gemineoa* eggs were less fertile than other crosses. Together, these findings argue that reproductive barriers accumulating across early life stages could explain the limited hybridisation of species in nature (Gallegos et al., 2023), providing new insights into the effects of temperature and sex on reproductive barriers, in present and future climates.

That reproductive barriers were strongest in larvae, when thermal niches differed most between species, suggests that niche differentiation could influence their isolation. Species’ ranges diverge along a latitudinal gradient noted for widespread clinal adaptation (Wood et al., 2021), and species show genomic signals of thermal adaptation in the hotspot (Gallegos et al., 2023). Such signals are now found to map to divergent thermal optima in purebred larvae, although *G. gemineoa* larvae had lower survival overall. This coincides with lower genetic diversity in *G. gemineoa* (Gallegos et al., 2023), but this link requires validation. Whether divergent adaptation helped species barriers to form is unknown, and genetic incompatibilities that make F1 hybrids inviable, or lead to hybrid breakdown in later generations, can emerge from either niche-driven divergence in sympatry or divergence in allopatry followed by secondary contact (Butlin & Smadja, 2018; Dobzhansky, 1937; Keller & Seehausen, 2012). Further work is therefore needed to disentangle these scenarios in *Galeolaria*. Adaptation aside, our results support the accumulation of incompatibilities in genes acting later in development (Orr & Turelli, 2001; Ortíz-Barrientos et al., 2007), which temperature may exacerbate by increasing biological rates and energy demands as life stages become more temperature-sensitive (Hochachka & Somero, 2002; Hoekstra et al., 2013). Although sensitivity initially increases at embryogenesis in *Galeolaria* and other external fertilizers (Dahlke et al., 2020; Rebolledo et al., 2020), lack of substantial barriers at this stage could reflect constraints on early-acting genes that make embryogenesis in the external environment particularly robust (Hamdoun & Epel, 2007). Ultimately, it will be important to test whether postzygotic barriers linked to thermal tolerance in hybrids continue to strengthen as development unfolds in *Galeolaria*, as seen in other systems (Álvarez & Garcia-Vazquez, 2011).

Parent-of-origin asymmetry in barriers at larval development further argues that maternally inherited factors control the thermal niches of hybrids. This could implicate the mitochondrial genome, given that mitochondria are usually inherited solely from mothers and play a crucial role in thermoregulation (Breton & Stewart, 2015; Hochachka & Somero 2002). Mitochondrial haplotypes are known to shape thermal niches (Chung & Schulte, 2020; Quintela et al., 2014) and vary latitudinally in association with climate (Camus et al., 2017; Silva et al., 2014). Further, functional or genetical incompatibilities with novel nuclear backgrounds can impose selection on hybrids in ways that interact with thermal niches of parental lineages (Ellison et al., 2008; Keller & Seehausen, 2012). It is therefore plausible that cytonuclear interactions involving mitochondria underpin the asymmetry of temperature-dependent barriers in *Galeolaria*. Alternatively, given the small size of the mitochondrial genome, other maternally-derived factors such as endosymbionts (Bruzzese et al., 2022; Gebiola et al., 2016), or sex-biased gene expression (Badyaev, 2013), could also play a role. Since asymmetry in reproductive isolation is widespread in nature (Haldane, 1922; Turelli & Moyle, 2007), but its mechanisms and links to temperature are unclear (e.g., Hudson et al., 2021; Martins et al., 2019), these gaps are ripe for study to better understand the stability of species barriers in a warming world.

Prezygotic barriers were also asymmetric, given that hybrid crosses with eggs (but not sperm) of *G. gemineoa* were less fertile than pure crosses (as also reported by Styan et al., 2008), but only weakly depended on temperature. Fertilization has a remarkably broad thermal niche in *Galeolaria* and other external fertilizers (Byrne et al., 2010; Rebolledo et al., 2020), so it is perhaps unsurprising that barriers at this stage were similarly robust. Nevertheless, sexual selection based on gamete interactions at fertilisation can drive gametic incompatibilities, and asymmetries in them, in this mating system (Kosman & Levitan, 2014; Martín-Coello et al., 2009). Such interactions, along with temperature, also mediate selection on sperm morphology in *Galeolaria* (Chirgwin et al., 2020; Johnson et al., 2013; Monro & Marshall, 2016), and we observed sister species’ sperm to be morphologically distinct (Figure S2). These lines of evidence, and new results here, thus suggest that the role of sexual selection in prezygotic barriers may be important to probe — for example, using competitive fertilisation assays to evaluate preferential fertilization of eggs by conspecific sperm. Moreover, that asymmetries in prezygotic *versus* postzygotic barriers responded differently to temperature points to differing mechanisms for their evolution, and highlights the importance of disentangling barriers at different life stages.

On a methodological note, marine species like *Galeolaria* present new opportunities, but also inherent challenges, for studying species barriers (Lotterhos et al., 2021; Marshall et al., 2016).

Despite evidence that the strength of postzygotic barriers may change through development (e.g., Álvarez & Garcia-Vazquez, 2011; Bundus et al., 2015), few studies have tested barriers across complete life cycles, and even fewer have tested barriers from one generation to the next, as necessary to infer hybrid breakdown (but see Hwang et al. 2016; Willett & Burton, 2003). This may speak to difficulties in rearing many species *in vitro*, as is the case for *Galeolaria* (Watson et al., 2017). Consequently, we could not trace offspring to maturity in this study. Moreover, we tested the effects of temperature at each life stage by standardizing thermal history at prior stages, as is common practice (Sinclair et al., 2016), but the potential for effects to aggregate over time (Levy et al., 2015; Williams et al., 2016) makes testing cumulative exposures a key next step. Last, we focused on extrinsic barriers controlled by temperature, given its key role in redistributing marine biodiversity (Lenoir et al., 2020; Sunday et al., 2015), but acknowledge that other variables (e.g., salinity) may also control barriers (Hwang et al., 2016).

Overall, our study presents new insights into the interplay of temperature and sex in reproductive barriers across early life stages, and points to shifting strengths of reproductive isolation in future climates. For example, if sympatry of *Galeolaria* species is relatively recent (given past support for allopatry; Halt et al., 2009; Styan et al., 2008), then climate change could already be shifting *G. gemineoa* poleward, and do so into the future (Sunday et al., 2015). On the one hand, this could see prezygotic barriers strengthen to limit gamete wastage as secondary contact intensifies (Coyne & Orr, 1989, 1997), or postzygotic barriers strengthen as pure offspring outperform hybrids at warmer temperatures. On the other hand, it could see postzygotic barriers weaken as maternal hybrids of *G. caespitosa* outperform pure offspring of *G. gemineoa* at cooler temperatures, and warm-adapted alleles introgress into *G. caespitosa* (Hamilton & Miller, 2016; Owens & Samuk, 2020). Such predictions need testing (e.g., with crosses of sympatric *versus* allopatric populations from contrasting thermal environments), but highlight the importance of assessing reproductive barriers and hybrid viability across ecologically-relevant environments to further our understanding of reproductive isolation, its fate, and the repercussions for biodiversity, in a rapidly-changing climate.

## Acknowledgements

We thank Emily Belcher for valuable help with laboratory work, Javiera Olivares for valuable help in sampling of specimens, and Fisheries Victoria (RP1328) and Parks Victoria (10008784) for collection permits. This project was supported by The Holsworth Wildlife Research Endowment & The Ecological Society of Australia awarded to CG, and by grants awarded under the Australian Research Council’s Discovery Scheme to KM and KH.

## Data availability statement

The data that supports the findings of this study, as well as reproducible R scripts with analyses and visualisations will be available on GitHub (https://github.com/CristobalGS/TemperatureSex-and-ReproductiveBarriers) upon publication.

