## Supplementary materials for "Temperature and sex shape reproductive barriers in a climate change hotspot"

**Table S1.** Sampling locations and dates.

| **Species** | **Location** | **Latitude** | **Longitude** | **Sampling date** |
| --- | --- | --- | --- | --- |
| *G. caespitosa* | Cape Otway | -38.85415 | 143.5485 | 4/11/2020 |
| *G. caespitosa* | Lorne | -38.54768 | 143.98691 | 28/04/2021 |
| *G. caespitosa* | Flinders | -38.47556 | 145.02573 | 29/04/2021 |
| *G. caespitosa* | YCW Beach | -38.50459 | 145.2514 | 23/06/2021 |
| *G. caespitosa* | Eagles Nest | -38.69295 | 145.70852 | 23/06/2021 |
| *G. caespitosa* | Walkerville | -38.86056 | 146.00061 | 8/11/2020 |
| *G. gemineoa* | Red Bluff | -37.86757 | 148.06232 | 7/11/2020 |
| *G. gemineoa* | Salmon Rocks | -37.80935 | 148.72623 | 25/04/2021 |
| *G. gemineoa* | Mallacoota | -37.57102 | 149.76412 | 6/11/2020 |
| *G. gemineoa* | Greenglades | -37.2824 | 149.94291 | 26/06/2021 |
| *G. gemineoa* | Merimbula | -36.89089 | 149.92981 | 26/06/2021 |
| *G. gemineoa* | Tathra | -36.72652 | 149.98494 | 24/04/2021 |

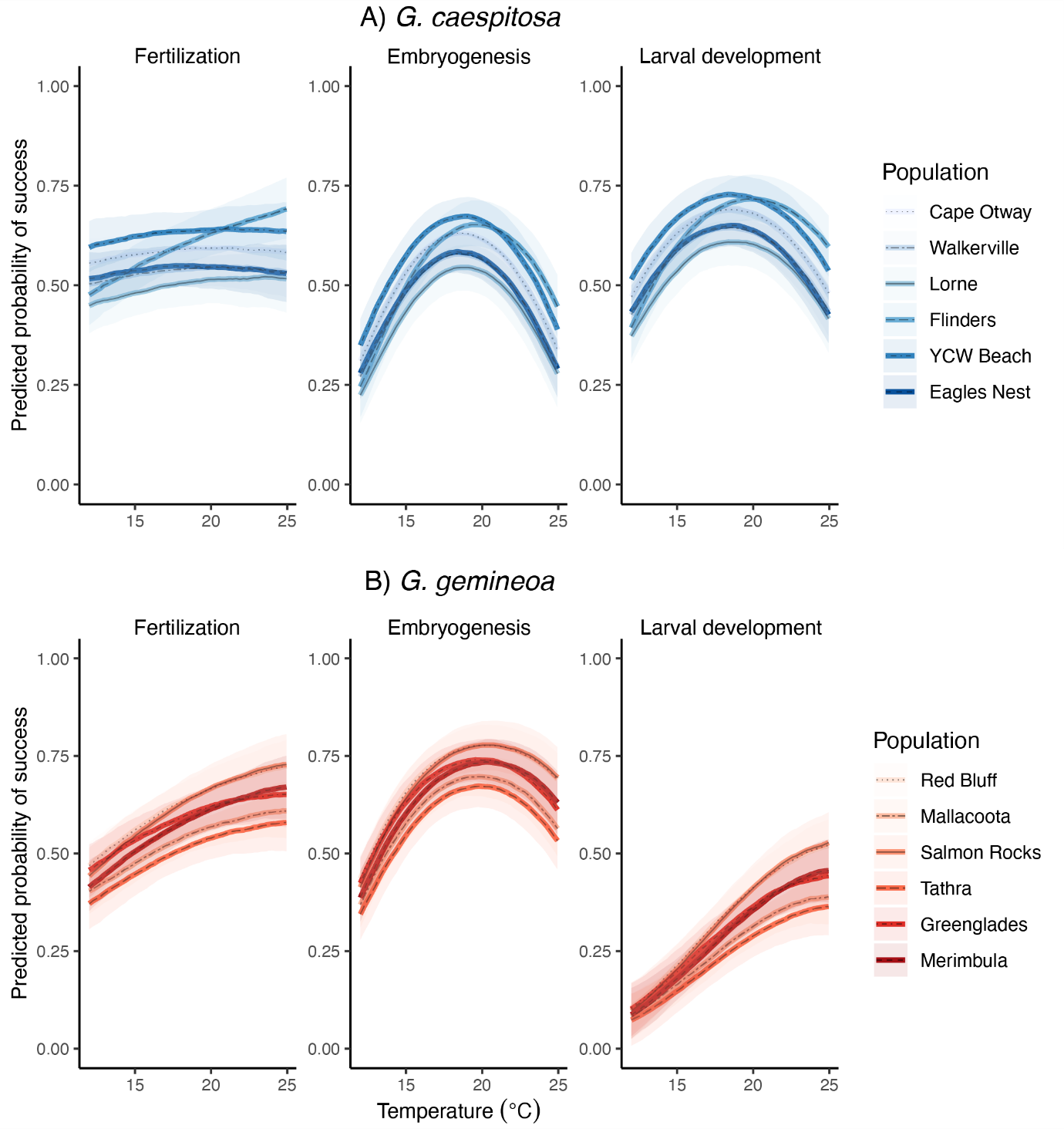

**Figure S1.** Thermal tolerance curves showing the predicted probabilities of fertilization success, survival of embryogenesis, and survival of larval development for populations of A) *G. caespitosa* (upper panels) and B) *G. gemineoa* (lower panels). Curves are predicted from population-level random effects estimated in a binomial mixed-effects regression of success on temperature interacting with cross and life stage (see main text). The legend for each species shows populations ordered by geographic location from southwest (top) to northeast (bottom). Shaded areas are 95% confidence intervals of curve predictions for random effects only, making variation among curves representative of population-level variation.

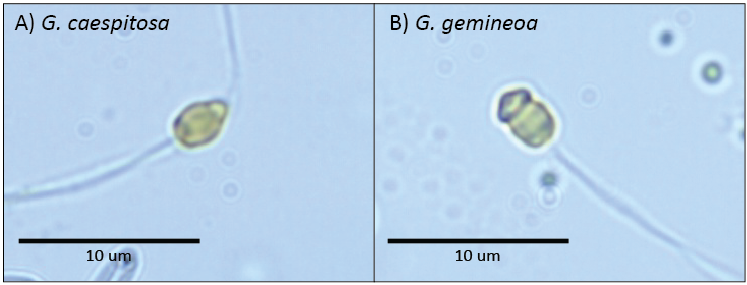

**Figure S2.** Typical sperm morphologies for A) *G. caespitosa* and B) *G. gemineoa*. Photos were taken using a digital camera mounted on a compound microscope at 100x magnification.
